# Cell surface ATP6V1B2 marks a subset of persistent senescent cells with increased resistance to apoptosis

**DOI:** 10.1101/2025.11.30.691415

**Authors:** Nataly Freizus, Julia Majewska, Yossi Ovadya, Ekaterina Kopitman, Ziv Porat, Avi Mayo, Tomer Meir-Salame, Bareket Dassa, Gil Stelzer, Uri Alon, Valery Krizhanovsky

## Abstract

Accumulation of senescent cells promotes ageing and age-related diseases. While senescent cells are heterogenous and increasingly persistent in vivo with age, the mechanisms underlying their heterogeneity, resistance to apoptosis, and tissue accumulation remain insufficiently understood. Here we report that in response to DNA damage, a subset of senescent cells upregulates the v-type ATPase subunit, ATP6V1B2 (V1B2) on the cell surface. This upregulation is associated with altered lysosomal activity and changes in intracellular pH. Heterogeneity of senescent cells marked by cell surface V1B2 (csV1B2) is present in naturally occurring senescent cells within both ageing and fibrotic lungs. Senescent cells expressing csV1B2 show an age-independent transcriptional signature associated with DNA repair and resistance to apoptosis. Consistent with this, we show that csV1B2 expression correlates with senescent cell resistance to ABT-737-induced apoptosis in culture. Our study identifies a subset of senescent cells, marked by csV1B2, with a distinct signature of apoptosis resistance. Understanding the functional heterogeneity of senescent cells and the mechanisms accountable for persistence of specific subpopulations in tissues may facilitate the development of improved senotherapeutic strategies for age-related diseases.

## Introduction

Cellular senescence is a state of stable growth arrest of metabolically active cells. It plays a pivotal role throughout life, starting as early as embryonic development where it governs organ remodeling^1,2^, and continuing as a tumor suppressor and an anti-fibrotic mechanism^3,4^. However, the progressive accumulation of senescent cells (SnCs) across tissues is a hallmark of aging and contributes to the onset and progression of age-related diseases, including cancer, ultimately limiting healthspan and lifespan^5,6^.

The physiological and pathological role of SnCs depends on their persistence in vivo, which is regulated by immune-mediated clearance, but also their intrinsic resistance to apoptosis^7,8^. Through secretion of cytokines, chemokines, and other factors collectively known as the senescence-associated secretory phenotype (SASP), SnCs recruit immune cells. Subsequent interactions between innate and adaptive immune cell receptors and SnC surface markers orchestrate their recognition and elimination. While surface expression of Vimentin, DPP4, MICA, and ULBP2 on SnCs facilitates their immune-mediated clearance, upregulation of PDL-1/2, GD3, and HLA-E enables their immune evasion ^9–15^. Thus, senescence surveillance depends on the effector functions of the immune system, with immunoregulatory markers on SnCs orchestrating either efficient or impaired clearance. Yet, whether next to immunosurveillance there are intrinsic mechanisms contributing to the heterogeneity SnCs accumulation in tissues and their apoptotic resistance remains to be elucidated.

SnCs are highly heterogeneous, depending on cell type, tissue microenvironment, and the nature of the inducing stressor^16^. This complexity presents a challenge as existing markers often lack specificity and fail to reflect the functional heterogeneity of SnCs. Therefore, identifying molecular markers that define functionally heterogenous SnC subpopulations, especially those linked to persistence and apoptosis resistance, is critical for understanding their role in ageing and disease.

To identify novel extracellular markers of SnCs we performed a proteomic screen for plasma membrane proteins enriched on SnCs. We found that DNA damaged-induced senescent fibroblasts upregulate the membrane-embedded subunit ATP6V1B2 (V1B2) of v-type ATPase proton pump. We observed cell surface expression of V1B2 on SnCs in ageing and fibrotic lung across multiple cell types. Notably, only a subset of SnCs exhibits surface expression of V1B2, suggesting heterogeneity within senescent population. Interestingly, sequencing of V1B2-membrane expressing senescent cells has revealed a conserved and age-independent transcriptional signature coupled to DNA damage and resistance to apoptosis. Consistent with this, we show that membrane expression of V1B2 is associated with resistance to ABT737-induced apoptosis in SnCs in vitro. Our findings suggest that the membrane expression of V1B2 in SnCs is associated with a selective DNA damage response mechanism that promotes resistance to apoptosis and may contribute to persistence of SnCs in ageing tissues.

## Results

### SnCs upregulate cell surface V1B2

Cellular senescence is associated with profound changes in the extracellular membrane proteome^17^. However, none of these changes specifically identify senescent cells (SnCs) or relate to their functional heterogeneity. To identify surface proteins associated with DNA damage induced SnCs, we performed a comparative proteomic screen for external plasma membrane–enriched fractions from proliferative and senescent cells (SnCs). We used a classical model of cellular senescence in primary human (IMR-90) lung fibroblasts induced to senesce by DNA damage^8^. Mass spectrometry-based peptide profiling of the ‘surfaceome’ across proliferating and SnCs revealed 1,954 differentially expressed proteins. To select proteins that were increased in senescence compared to corresponding proliferating control we applied stringent filtering criteria: a senescent-to-proliferating fold change greater than three, a minimum two-fold change in at least two independent replicates, sequence coverage exceeding 15%, and a p-value below 0.05. From 13 shortlisted proteins, 6 were prioritized for further experimental validation based on their biological function (ATP6V1B2, MXRA5, RPL10, SEPT2, MRP1, EPHB1; Fig. 1a-b). Compared to growing cells, SnCs exhibited significantly increased surface ATP6V1B2 (V1B2) expression across multiple human cell lines, including primary bronchial/tracheal epithelial cells, BJ fibroblasts and IMR-90 fibroblasts (Fig. 1c and data not shown). Notably, V1B2 was the only protein to exhibit such distinct and consistent surface labeling of SnCs across all cell lines tested. Furthermore, imaging flow cytometry analysis revealed a distinct membrane staining of V1B2 in senescent but not in growing IMR90 cells (Fig. 1d). Therefore, we identified V1B2 as a potential cell surface marker of DNA-damage induced SnCs.

**Figure 1.**
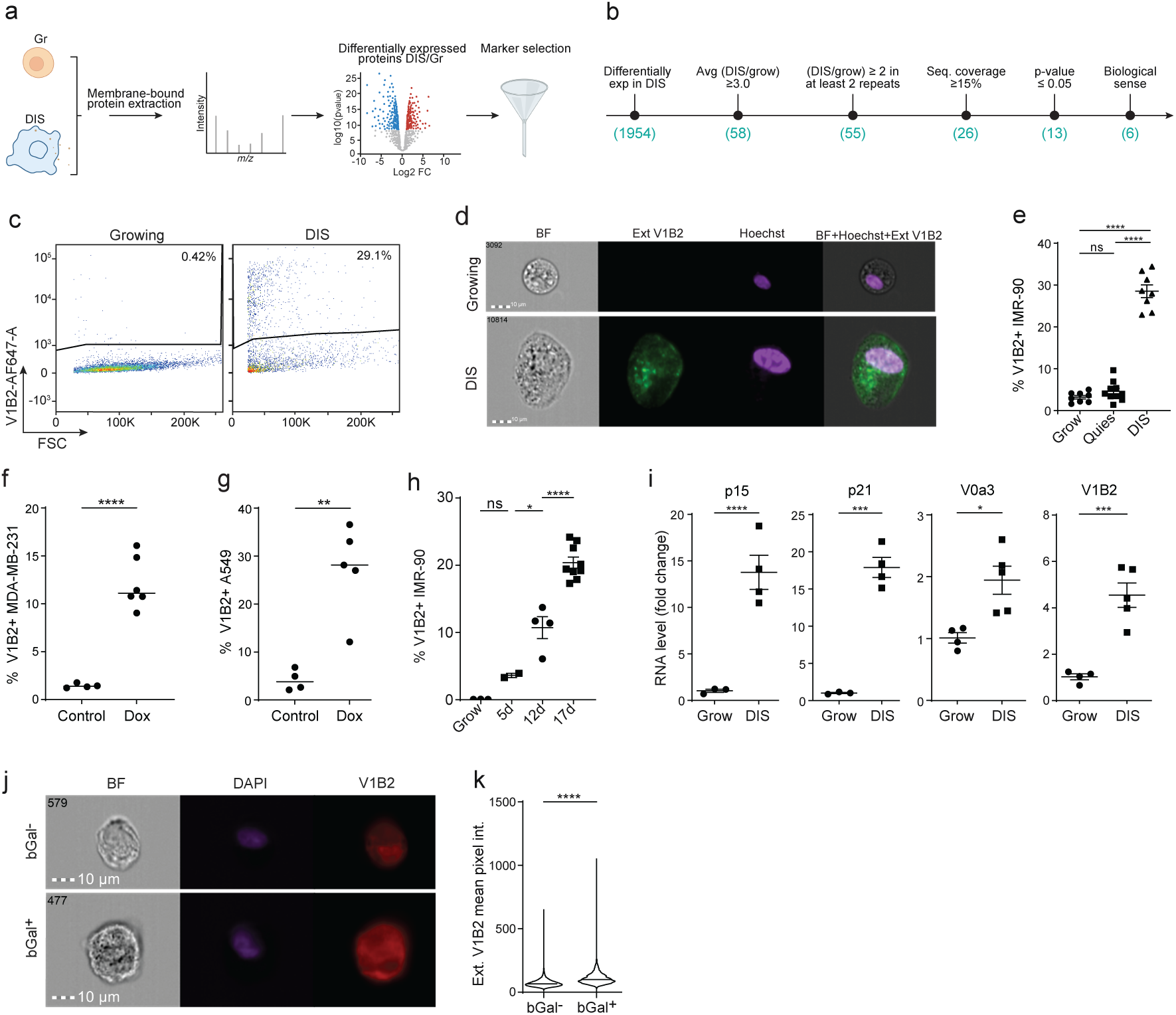
Senescent cells express V1B2 on cell surface. **a,** Experimental outline: external plasma membrane bound proteins from growing and DNA damage induced senescence (DIS) IMR-90 fibroblasts were extracted, identified by MS and analyzed for differential expression. **b,** Screening parameters for identification of differentially expressed cell surface proteins. **c,** Flow cytometry (FC) plots of csV1B2 vs. forward scatter (FSC) growing and DIS IMR-90 fibroblasts. **d,** Imaging FC of selected examples of extracellular V1B2 (ext V1B2) staining in live growing and DIS IMR-90 fibroblasts. The images of Bright Field (BF), ext V1B2 (green), nuclear staining labeled by Hoechst and their combination are presented. **e,** Percentage of V1B2+ cells in growing, quiescent and DIS IMR-90 fibroblasts. **f,** Percentage of V1B2+ cells in growing, and doxorubicin treated (DOX) MDA-MB-231 and (**g**) A549 cancer cell lines. **h,** Percentage of V1B2+ cells in growing and DIS IMR-90 fibroblasts at 5-, 12- and 17-days post senescence induction. **i,** Relative RNA expression of p15, p21, V0a3 and V1B2 in growing and DIS IMR-90 fibroblasts. **j-k,** Imaging FC and quantification of idiopathic pulmonary fibrosis patient-derived lung fibroblasts stained for SA-β-galactosidase activity (bGal), and for ext V1B2 expression. The bGal is visualized by dark staining in BF. **k**, Ext V1B2 expression was quantified in bGal positive (bGal+) and bGal negative (bGal-) cells. Two tailed t-test was used for pairwise statistical comparisons; ordinary one-way ANOVA test was used for multiple statistical comparisons. The data are presented as mean ± SEM; *p<0.05, **p<0.01, ***p<0.001, ****p<0.0001. Quantified results represent at least three experimental repeats.

Senescence and quiescence are two distinct cellular states characterized by cell cycle arrest, but they have fundamentally different biological functions^6,18^. To understand if V1B2 surface expression is linked to proliferative arrest, we evaluated V1B2 surface levels in both senescent and quiescent IMR-90 fibroblasts. Our results show a significant increase in surface V1B2 level only in SnCs, compared to both growing and quiescent cells, with 25% of SnCs staining positive for cell surface V1B2 (csV1B2) expression (Fig. 1e). Interestingly, we also found that senescent cancer cells exhibit over a 6-fold increase in csV1B2 level relative to untreated cancer cells observed in both MDA-MB-231 (p<0.0001) and A549 cells (p<0.05; Fig. 1f-g). These results suggest that V1B2 surface expression is not merely a feature related to cell cycle arrest, but is selectively upregulated in senescence, regardless of cells initial proliferative potential or the cell type.

Senescence is a dynamic cellular state characterized by temporal changes in gene expression pattern, reflecting its evolving cellular phenotype^19,20^. To understand if V1B2 surface expression changes over time, we measured csV1B2 level in senescent IMR-90 fibroblasts at 5-, 12-, and 17-days post senescence induction. We observed a gradual increase in the percentage of V1B2 surface staining in SnCs, with approximately 25% of SnCs exhibiting surface V1B2 staining by day 17 (Fig. 1h). These results suggest that the DNA damage and potentially downstream repair mechanisms may regulate the level of membrane V1B2 expression in senescent cells.

V-ATPases are complexes composed of fourteen subunits, eight clustered into a peripheral V1 domain responsible for ATP hydrolysis, and six clustered into a membrane integral V0 domain. V-ATPase function is regulated by the isoform’s variability across its subunits. Among these, the four isoforms of the V0a subunit are particularly well described. V0a1 and V0a2 anchor V-ATPases to synaptic vesicles and early endosomes, respectively, while V0a3 and V0a4 regulate translocation to the plasma membrane in diverse cell types^21,22^. To understand regulation of the subcellular localization of V1B2 in cellular senescence, we checked the expression of the ATP-ase domains, together with senescence markers, in growing and senescent IMR-90 fibroblasts. Along with elevation in senescence markers p15 and p21, SnCs also upregulated V0a3 and V1B2 expression (Fig. 1i). The upregulation of V0a3 expression supports the potential re-localization of V-ATPase to the plasma membrane in senescent cells.

SnCs are present in different age-related pathologies. We, therefore, explored membrane expression of V1B2 in a clinically relevant in-vitro senescence model, lung fibroblasts from idiopathic pulmonary fibrosis (IPF) patients. To identify SnCs we stained cells for β-galactosidase activity (βGal) ^23^. Subsequently, cells were stained for V1B2 and analyzed using imaging flow cytometry. In this experimental setting, whole-cell V1B2 content was stained, and external expression was quantified by the fluorescent signal on the periphery of cell area. We observed a significant elevation in V1B2 membrane level in βGal+ cells, relative to βGal- cells (p<0.001, Fig. 1j-k). Overall, these results suggest that cell surface expression of V1B2 is upregulated in SnCs across cell types in physiological and pathological conditions.

### CsV1B2 marks senescent cell functional heterogeneity

Heterogenous expression of csV1B2 in senescent cells prompted us to evaluate the functional significance of V1B2-positive (V1B2+) compared to V1B2-negative (V1B2-) cells by transcriptomic profiling. Principal component analysis (PCA) revealed distinct separation based on V1B2 surface expression, with PC1 and PC2 explaining 31.6% and 13.8% of the variance, respectively (Fig. 2a). V1B2+ cells showed a significantly more upregulated than downregulated differentially expressed genes (DEGs). A global transcriptional activation may mark senescent cell adaptation to stress. Notably, several genes encoding V-ATPase subunits – including ATP6V1AP2, ATP6V1A, and ATP6V1B2 (encoding V1B2 subunit itself) were upregulated in V1B2+ cells (Fig. 2b). Additionally, genes associated with cell death pathways (BIRC2, BNIP3L, BNIP2, API) were also significantly enriched in V1B2+ cells (Fig. 2b). Indeed, pathway enrichment analysis of differentially expressed genes (DEGs) revealed pathways that relate to cell death, including regulation of TP53 activity, P70S6K signaling, programmed cell death, mTOR signaling and intrinsic pathways for apoptosis (Fig. 2c).

**Figure 2.**
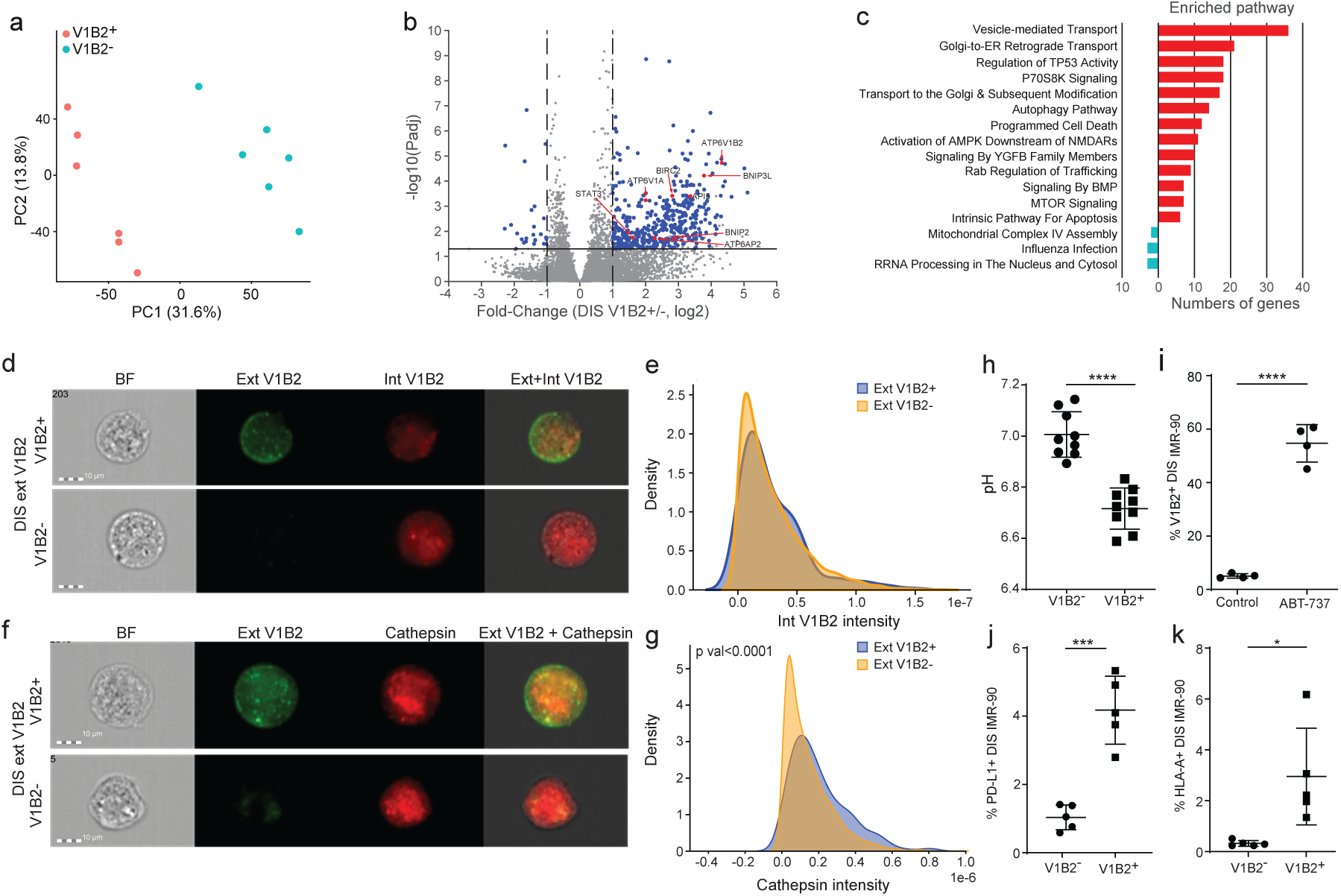
V1B2+ senescent cells enhance lysosomal activity and resistance to apoptosis. **a,** Principal component (PC) analysis of V1B2- and V1B2+ DIS IMR-90 cells. **b,** Volcano plot of 466 differentially expressed genes (DEGs) between V1B2+ and V1B2- DIS IMR-90 cells. DEGs were identified by Padj <= 0.05 and |log2 Fold change | >= 1. ATP6V1B2 and apoptotic signaling-related genes are labeled. **c.** Pathway enrichment analysis of DEGs identified in (b). Enriched super-pathways are ordered by number of genes identified in each pathway. Enrichment in V1B2+ cells (red), enrichment in V1B2- cells (green). **d-e,** Imaging FC images (d) and histograms (e) of DIS IMR-90 fibroblasts stained for external V1B2 (Ext V1B2), and internal V1B2 (Int V1B2). **f-g,** Imaging FC images (f) and histograms (g) of DIS IMR-90 cells stained for Ext V1B2 and Cathepsin B. **h,** Intracellular pH measurement of V1B2+ and V1B2- DIS IMR-90 **i.** IMR-90 fibroblasts were treated with 10uM ABT-737 or vehicle (control) and the percentage of V1B2+ cells following the treatment is presented. **j-k,** Percentage of PD-L1 (j) and HLA-A (k) on V1B2+ and V1B2-DIS IMR-90 fibroblasts. Two tailed t-test was used for pairwise statistical comparisons: *p<0.05, ***p<0.001, ****p<0.0001. Results represent at least three experimental repeats.

Another category of the top enriched pathways relates to vesicular transport (Golgi to ER retrograde transport, transport to the Golgi and subsequent modifications, autophagy pathways) (Fig. 2c). Given that V1B2 is physiologically localized to endo-membranes associated with vesicular transport, we asked whether its increased surface level could result from elevated intracellular V1B2 levels and enhanced trafficking to the plasma membrane. To this end, we did extracellular staining for V1B2 on live senescent IMR-90 fibroblasts, followed by fixation/permeabilization and additional staining to detect total intracellular V1B2 levels. Based on computational masks that segment the cell surface area into peripheral (20%) and internal (80%) regions we could distinguish spatial localization of V1B2 staining. There was no significant difference in intracellular V1B2 levels between cells with extracellular V1B2 staining and their negative counterparts (Fig. 2d-e). These results suggest that the increased V1B2 membrane level is not dependent on intracellular V1B2 protein level but rather may result from its increased trafficking to the membrane.

Lysosomal dysfunction is extensively linked to cellular senescence through elevation of lysosomal enzyme activity (SA-b-galactosidase)^24^, lysosomal accumulation of lipofuscin^25^, lysosomal expansion, lysosomal membrane permeabilization and pH alterations^26^. Under physiological conditions, V1B2 localizes to the lysosomal membrane, where it functions as a component of the ATP-ase complex regulating lysosomal pH^27^. However, the functional significance of extracellular V1B2 in regulation of lysosomal activity and intracellular pH, remains unknown. To address this question, we investigated lysosomal activity by examining cathepsin levels in IMR-90 cells expressing extracellular V1B2. Surface expression of V1B2 correlated with elevated levels of cathepsin, suggesting an increase in overall lysosomal content or lysosomal activity (Fig. 2f-g). Moreover, cells with membrane V1B2 expression showed a significant decrease of intracellular pH (p<0.0001, Fig. 2h), supporting a possibility of an active role of extracellular V1B2 subunit in proton transport. Altogether these results suggest a possible functional role of membrane V1B2 as part of ATP-ase complex in the regulation of lysosomal activity and intracellular pH.

SnCs upregulate anti-apoptotic BCL-2 family proteins, which support their apoptosis resistance and their persistence within tissues^8^. Sequencing of csV1B2-membrane expressing SnCs has revealed a transcriptional signature coupled to intrinsic apoptosis pathways. This suggested that V1B2 cell surface expression might be correlated with the regulation of apoptosis of these cells. To address this correlation, we treated senescent IMR-90 fibroblasts with a senolytic drug ABT-737, which inhibits BCL-2 family proteins to selectively induce apoptosis in these cells. Strikingly, we found an eleven-fold increase in the percentage of csV1B2+ cells following the ABT-737 treatment (p<0.0001, Fig. 2i), suggesting that this subpopulation is more resistant to apoptosis induction. Additionally, cells expressing csV1B2 showed a significant upregulation of PD-L1 and HLA-A, proteins involved in immune evasion mechanisms of SnCs (Fig. 2j-k) ^12,14,15,17,28^. Altogether, these results suggest that csV1B2+ cells may employ both intrinsic apoptosis resistance and immune evasion mechanisms facilitating their persistence in tissues.

### The presence of csV1B2 correlates with p16 expression in vivo

The cell surface expression of V1B2 marks a subpopulation of SnCs in vitro. Thus, we asked if there is a similar heterogeneity of cell surface presence of V1B2 on senescent cells *in vivo*. To this end, we used a mouse model of bleomycin-induced lung fibrosis, as a model in which the contribution of senescence to the disease pathology is well-established^29^. Two weeks after injection of bleomycin or PBS, we analyzed lung epithelial cells expressing EpCam (EpCam+) for surface V1B2 expression using imaging flow cytometry following senescence-associated beta-galactosidase (bGal) staining. (Fig. 3a). As expected, the percentage of bGal+ epithelial cells was significantly increased in bleomycin-treated mice compared to those treated with PBS (Fig. 3b) ^30^. Whole-cell expression of V1B2 in either bGal- or bGal+ epithelial cells did not show a difference between PBS and bleomycin-treated mice (Fig. 3c-d). However, percentage of cells with surface expression of V1B2 was significantly, more than two-fold, increased only within bGal⁺ epithelial cells and not in bGal- epithelial cells when bleomycin-treated mice were compared to the PBS control mice. (p<0.05, Fig. 3e-g). Moreover, only a subset of about 40% of bGal+ cells upregulated extracellular expression of V1B2 consistent with our *in vitro* results. Therefore, the expression of V1B2 on the cell surface of senescent epithelial cells is increased *in vivo* in a lung fibrosis model, though its expression is heterogonous among SnCs.

**Figure 3.**
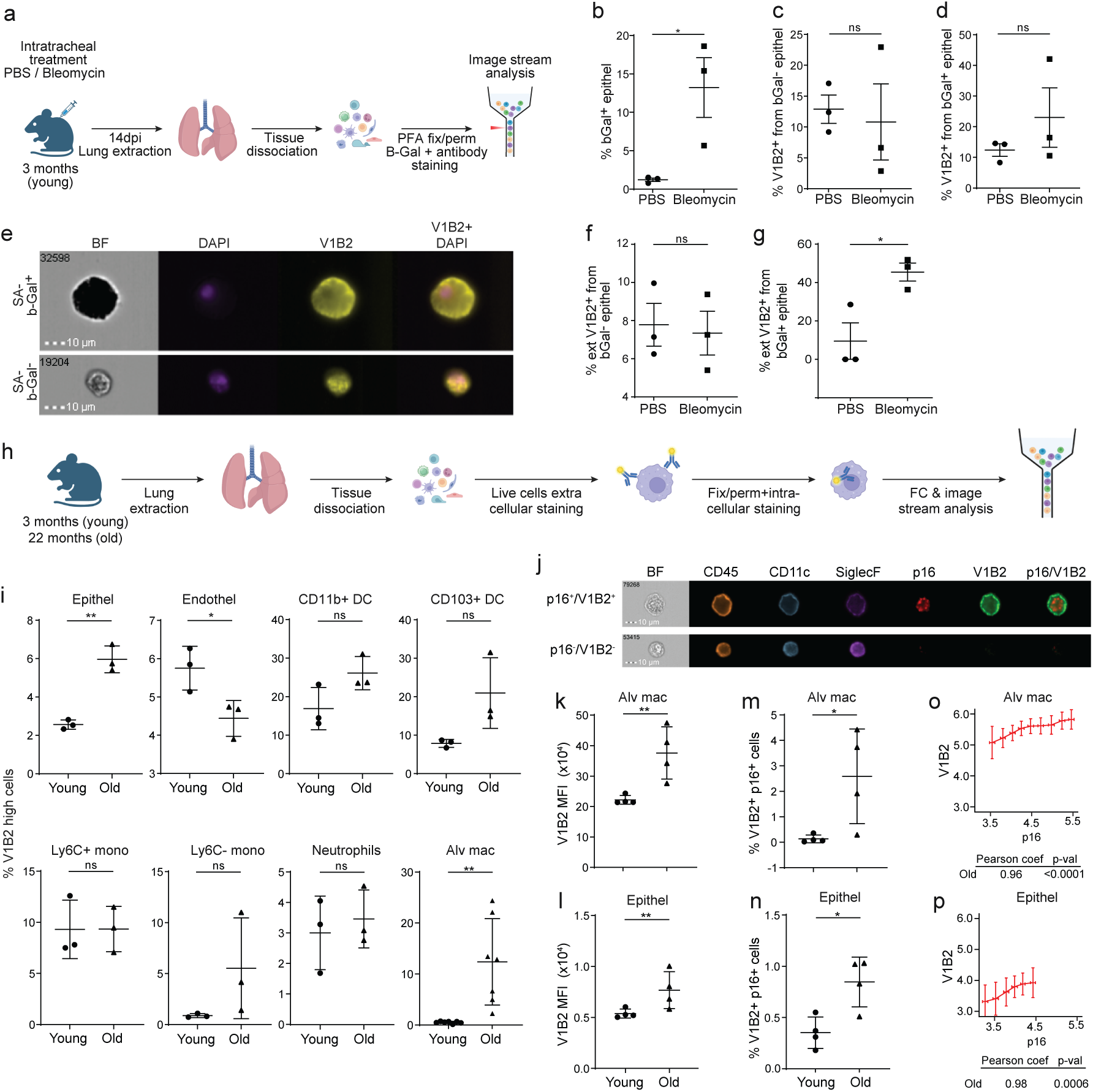
Cell surface expression of V1B2 correlates with p16 expression in-vivo. **a,** Experimental outline for evaluation of csV1B2 expression in a lung fibrosis model: 3 months old mice were treated intratracheally with Bleomycin or PBS. 14 days post treatment lungs were dissociated to single cells suspension, fixed, permeabilized and stained for SA-b-galactosidase activity. The cells were then, stained with DAPI and antibodies to CD45, EpCam and V1B2. **b,** Percentage of epithelial SA-b-galactosidase positive (bGal+) cells in PBS and Bleomycin treated mice. **c-d,** Percentage of total V1B2+ cells out of bGal- and bGal+ epithelial cells from PBS and Bleomycin treated mice. **e,** Imaging Flow Cytometry of bGal+ and bGal- epithelial cells. **f-g,** Percentage of external V1B2+ (ExtV1B2) cells within bGal- and bGal+ cells in PBS and Bleomycin treated mice. **h,** Experimental outline for evaluation of csV1B2 expression in the old (20-22 months old) and young (3 months old) mice. i, Percentage of V1B2+ cells in the indicated cell populations. **j,** Imaging FC of ExtV1B2+ p16+ double positive (p16+/V1B2+) and ExtV1B2-p16-double negative (p16-/V1B2-) alveolar macrophages stained with the indicated markers. **k-l,** Mean fluorescence intensity (MFI) of ExtV1B2 staining in alveolar macrophages (Alv mac) and epithelial cells (Epithel) derived from young and old mice. **m-n,** Percentage of ExtV1B2+ p16+ double positive (V1B2+p16+) alveolar macrophages (Alv mac) and epithelial cells derived from young and old mice. **o-p,** Pearson correlation between ExtV1B2 (V1B2) and p16 protein expression in alveolar macrophages and epithelial cells in old mice. The AM cells were binned into 9 bins in with a median of 4000 cells per bin, and epithelial cells were binned into 6 bins with a median of 4000 cells per bin. Single cells were ranked by p16 expression level in bins from low to high. For each bin, the mean expression level of V1B2 is shown. Pearson correlation coefficient and associated *P* value (*p-val*) is indicated below the graph. Two tailed t-test was used for pairwise statistical comparisons. Error bars are mean ± SEM; *p<0.05, **p<0.01.

(SnCs) not only contribute to a variety of age-related diseases but also drive tissue ageing through their accumulation^29,31–34^. However, it is unknow whether senescent cells with csV1B2 expression are persistent in ageing tissues. We therefore assessed expression of csV1B2 in combination with the intracellular senescence marker p16 in aged (20-22 months old) and young (3 months old) mice (Fig. 3h). We analyzed lung single cell suspension from these mice and focused on epithelial cells, endothelial cells and six subsets of immune cell populations, including dendritic cells (DC), monocytes, neutrophils and alveolar macrophages. Out of the above populations only epithelial cells and alveolar macrophages showed significant increase in the percentage of V1B2+ cells in the old mice compared to the young controls (Fig. 3i). We then analyzed alveolar macrophages for expression of p16 and V1B2 by imaging flow cytometry (Fig.3j) We observed p16 positive alveolar macrophages that exhibited a clear ring of V1B2 staining on the cell’s periphery (Fig.3j). We then compared mean fluorescence intensities (MFI) of V1B2 in both alveolar macrophages and epithelial cells (Fig. 3k-l). Consistent with the analysis of V1B2+ cell frequencies, the MFI-based analysis revealed an age-dependent increase in V1B2 expression, with 40% and 100% increase in MFI in epithelium and alveolar macrophages, respectively (p<0.01 and p<0.01, Fig. 3k-l). Therefore, cell surface expression of V1B2 is increased on epithelial cells and alveolar macrophages with age.

Senescent cells are identified by multiple markers, yet p16 shows the strongest and most consistent correlation with ageing and age-related diseases. Therefore, we were interested in studying the co-expression patterns of p16 and csV1B2. We focused on alveolar macrophages and epithelial cells in which csV1B2 expression is increased with age. The percentage of p16 and csV1B2 double positive cells was significantly higher in old mice compared to young mice for both alveolar macrophages and epithelial cells (p<0.05 and p<0.05, Fig 3m-n). Moreover, single-cell analysis of AM and epithelial cells revealed a positive correlation between p16 and V1B2 (Spearman’s rank correlation coefficient greater than 0.95 for both cell types; p<0.0001 and p<0.001, Figure 3o-p), These findings might suggest a link between core senescence pathways and surface expression of V1B2.

### Myeloid-biased V1B2 expression is linked to an age-independent response to DNA damage

Cell surface expression of V1B2 is increased with age in the selected cell populations. To advance our understanding of the nature and the molecular characteristics of V1B2+ cells in aged lung tissue, we performed single-cell RNA sequencing (scRNA-seq) on approximately 50,000 V1B2+ and V1B2- immune cells sorted by fluorescence-activated cell sorting (FACS) from young 2-month-old and aged 24-month-old C57BL/6 wild-type mice (Fig. 4a and Extended Data Fig. 4a). Using unsupervised clustering of CD45⁺ immune cells, combined with expression of canonical marker genes, we identified the major lung myeloid and lymphoid immune cell types, including monocytes (Mon), macrophages (Mac), alveolar macrophages (AM), interstitial macrophages (IM), dendritic cells (DCs), eosinophils (Eos), neutrophils (Neu), natural killer (NK) cells, B cells, CD4⁺ T cells, and cytotoxic CD8⁺ T cells (Fig. 4b and Extended Data Fig. 4b-c). Senescent cells, identified based on the SenMayo transcriptomic signature^35^, were predominantly found within the myeloid lineage, which showed markedly higher expression of senescence-associated genes compared to the lymphoid lineage (Fig. 4c). Moreover, V1B2+ cells were found predominantly within the myeloid lineage (Fig. 4d and Extended Data Fig. 4d-e).

**Figure 4.**
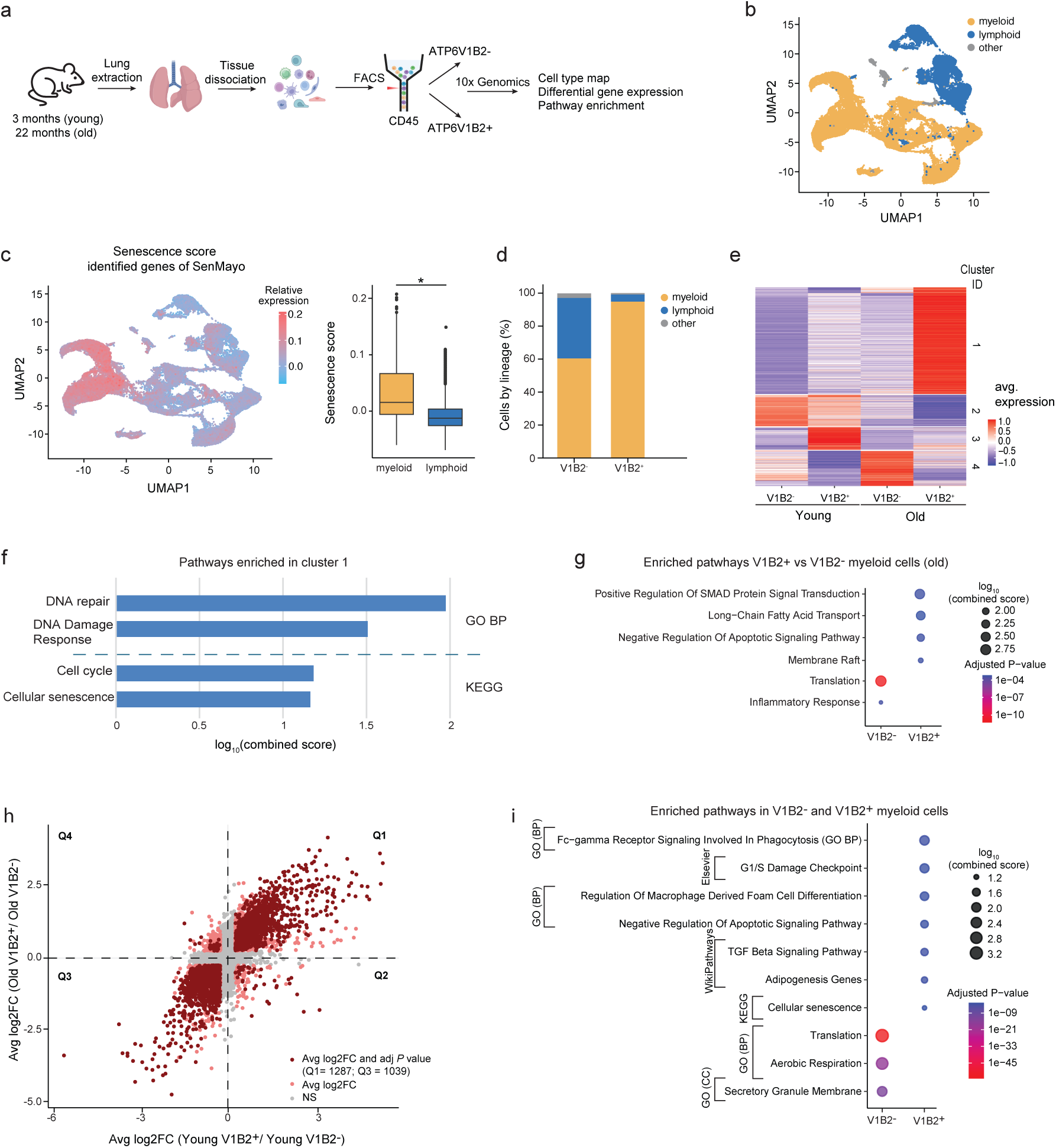
Myeloid-biased V1B2 expression is linked to an age-independent response to DNA damage. **a,** Experimental setup included lung single cell suspension from young and old mice followed by FACS-based sorting of V1B2-negative (V1B2-) and V1B2-positive (V1B2+) cells from CD45+ immune cells for scRNAseq analysis. **b,** Uniform manifold approximation and projection (UMAP) visualization of immune cell lineages (myeloid, lymphoid) based on identified immune cell populations. Colors represent different lineages as indicated. **c,** Module score of senescence signature in each cell, based on SenMayo gene list for mice^35^, is indicated by the color scale (left). Box plots representing quantification of senescence signature in indicated lineages (myeloid, n=38261 cells; lymphoid, n=13986 cells) (right). **d,** Bar plots showing the distribution of immune lineages in V1B2- and V1B2+ cells. **e,** Heatmap depicting distinct gene modules with similar expression patterns across V1B2- and V1B2+ cells of myeloid lineage from young and old mice. Color scale indicates scaled gene expression. **f,** Pathways significantly enriched in cluster 1 (e) based on KEGG and GO databases **g**, Pathways significantly enriched in V1B2+ versus V1B2- myeloid cells from old mice, based on Gene Ontology (GO) analysis. All pathways are derived from the GO Biological Process category, except for ‘membrane raft’, which is from GO Cellular Component category. **h,** Scatter plot showing average log2 fold change (Avg log2FC) of gene expression between V1B2+ and V1B2- myeloid cells in young (x-axis) and old (y-axis) mice. Each point represents a gene. Genes in Q1 are upregulated in V1B2+ cells in both young and old myeloid cells, while genes in Q3 are upregulated in V1B2- myeloid cells across age groups. Quadrants are defined by average log2FC threshold of 0. Avg log2FC=>0.25 and adj P value=<0.05. **i,** Pathways significantly enriched in V1B2+ cells (Q1) and V1B2- cells (Q3) in young and old mice (based on h). Pathways are derived from the following databases: Gene Ontology (GO) Biological Process (BP) and Cellular Component (CC) categories, as well as KEGG, Elsevier Pathway Collection, and WikiPathways, as indicated. Data are presented as mean ± s.e.m.; Statistical significance was assessed using two-sided Wilcoxon rank sum test (Mann–Whitney *U* test) (c, f, g, h, i). Only pathways with p adj value <0.05 are shown, enrichment is shown as a combined score, as previously described^56^. \**P* < 0.05, \*\**P* < 0.01, \*\*\**P* < 0.001. Young (*n* = 2) and old (*n* = 2) mice were used (a–i).

Based on enrichment of senescence-associated genes, the cell surface expression of V1B2 and the physiological significance of senescence in the myeloid populations in the lung ^14^, we decided to concentrate on identified myeloid cell types for subsequent transcriptomic analyses. To understand how transcriptional profile of myeloid cells differ with V1B2 expression and age, we performed principal component analysis. PCA showed distinct clustering of the four groups (Young V1B2^-^, Young V1B2^+^, Old V1B2⁻, Old V1B2⁺), with samples clustering primarily by V1B2 membrane expression, followed by age (Extended Data Fig. 4f). Further, k-means clustering allowed to differentiate distinct gene expression patterns (cluster 1, 2, 3 and 4) of V1B2-negative and V1B2-positive cells from young and old mice (Fig. 4e). Pathway enrichment analysis of genes upregulated in the cells positive for membrane V1B2 expression in old mice (cluster 1) indicated increased expression of DNA damage and repair, cell cycle checkpoints and cellular senescence pathways (Fig. 4f). Further analysis of V1B2+ and V1B2- myeloid cells from aged mice revealed a selective upregulation of apoptosis resistance related genes in V1B2+ population, as identified by the increase in the “negative regulation of apoptotic signaling pathway” (Fig.4g), and consistent with our previous findings (Fig.2c, 2i). Moreover, compared to V1B2-, V1B2+ myeloid cells were significantly enriched in gene signatures linked to SMAD signaling, membrane raft and lipid metabolism (Fig. 4g). Notably, V1B2+ cells showed reduced immune activation and translation, the latter being previously linked to senescence-associated transcriptional changes (Fig. 4g) ^36,37^. These findings suggest that membrane localization of the V1B2 in myeloid cells might coincide with an adaptive response to DNA damage and cellular stress such as senescence, associated with a globally reprogrammed state characterized by negative regulation of apoptosis, protein translation and altered immune function.

To understand whether the transcriptional signature of V1B2+ myeloid cells is conserved in ageing, we compared the gene expression fold change between V1B2+ and V1B2- cells in both young and old myeloid populations (Fig. 4h). This analysis identified 1287 genes which are consistently upregulated in V1B2+ cells in both age groups (Q1), while 1039 genes were consistently downregulated (Q3). Most of the differentially expressed genes followed similar directional changes in both age groups, suggesting that V1B2-associated gene expression is largely age-independent (Fig. 4h). Interestingly, genes upregulated in V1B2+ cells (Q1) were primarily enriched in pathways related to cell cycle, DNA damage checkpoints, senescence, regulation of apoptosis, TGF-B pathways and lipid metabolism (Fig. 4i). In contrast, genes upregulated in V1B2- cells (Q3) were associated with translation, aerobic respiration, and granule-mediated secretion (Fig. 4i). Overall, these results suggest that gene expression of V1B2-positive cells might be associated with specific phenotypic fitness traits such as DNA repair mechanisms in response to DNA damage, regardless of age or the microenvironment these cells reside in.

### CsV1B2 is associated with resistance to apoptosis in senescent myeloid cells in vivo

Senescent myeloid cells expressing csV1B2 exhibited marked resistance to apoptosis, which may contribute to their persistence in aged tissues. To better understand the functional contribution of these cells we examined the transcriptional profiles of csV1B2+ immune populations in aging. First, the proportion of csV1B2+ cells within the myeloid lineage was higher in old mice compared to young counterparts (Fig. 5a). Notably, V1B2+ cells were mostly present in macrophages (Mac I, AMs, IMs), dendritic cells (DCs) and eosinophils (Eos), pronounced in old mice compared to young but did not depend on overall relative abundance of these populations with ageing (Extended Data Fig. 5a-c). Moreover, expression of senescence-related genes correlated with cell surface localization of V1B2 (Fig. 5b). Additionally, csV1B2 was also associated with significantly higher resistance to apoptosis across these populations, except for eosinophils (Fig. 5c). Of note, we observed a strong positive correlation between resistance to apoptosis and the percentage of myeloid immune cell types with cell surface localization of V1B2, in both young (r^2^=0.7) and old (r^2^=0.76) mice (Fig. 5d). This finding suggests that cell surface localization of V1B2 might be a consequence of stress-response mechanisms that enhance resistance to apoptosis.

**Figure 5.**
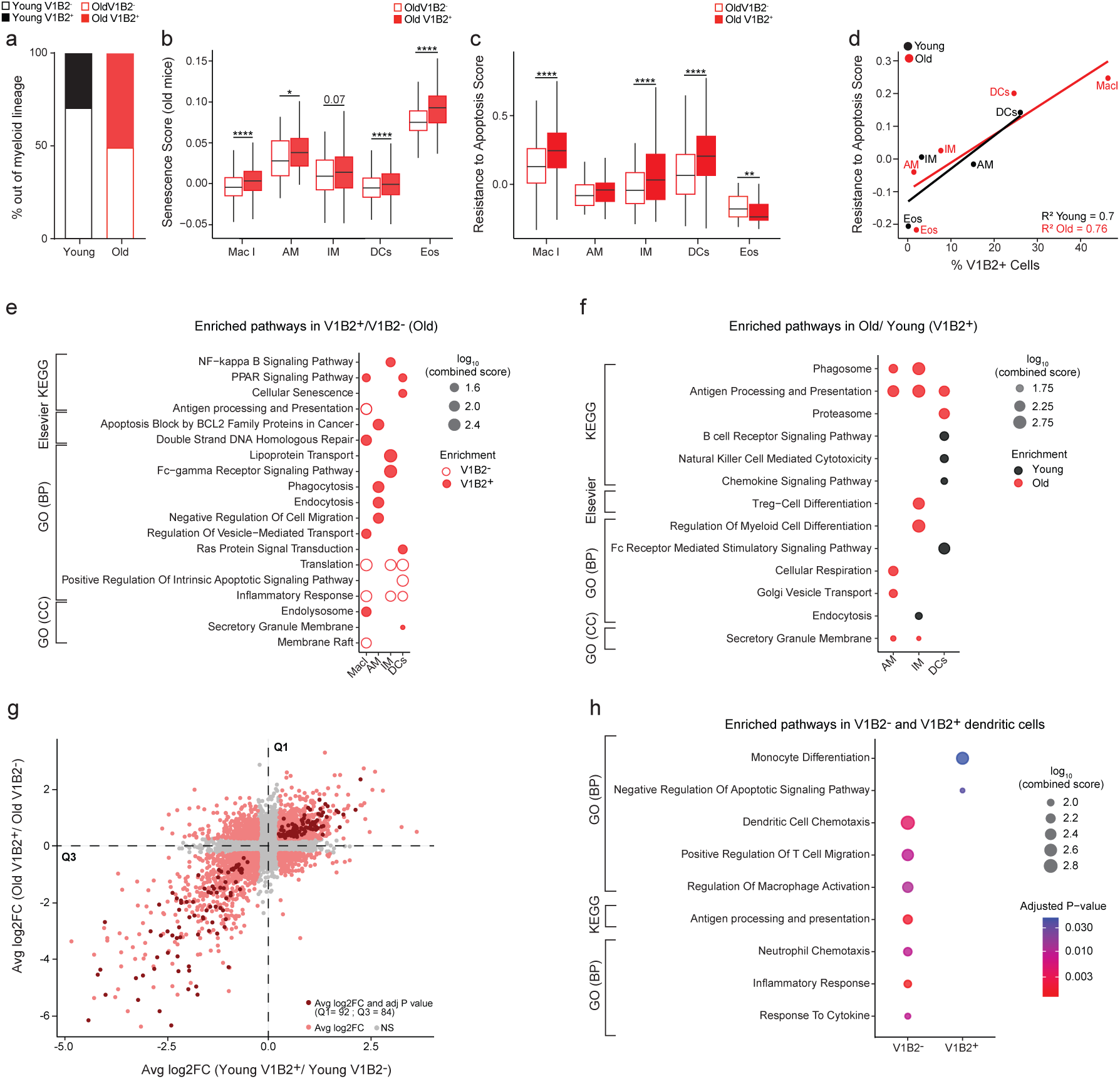
V1B2 expression is associated with senescence and resistance to apoptosis *in vivo*. **a,** Distribution of V1B2- and V1B2+ myeloid cells within young and old mice. **b,** Quantification of senescence-associated gene signature in V1B2- and V1B2+ immune cell types in old mice, as described previously^35^. **c,** Quantification of resistance to apoptosis-associated gene signature in V1B2- and V1B2+ immune cell types in old mice. **d,** Scatter plot showing the correlation between percentage of V1B2+ cells (x-axis) and resistance to apoptosis score (y-axis) across selected immune cell types in young (black) and old (red) mice. Linear regression lines are shown for each age group, with coefficient of determination (R^2^). **e,** Pathways significantly enriched in V1B2+ vs V1B2- cells in MacI, AM, IM, and DCs in old mice based on KEGG, Elsevier and GO databases. **f,** Pathways significantly enriched in V1B2+ old vs V1B2+ young cells in AM, IM and DCs based on KEGG, Elsevier and GO databases. **g,** Scatter plot showing average log2 fold change (Avg log2FC) of gene expression between V1B2+ and V1B2- DCs in young (x-axis) and old (y-axis) mice. Each point represents a gene. Genes in Q1 are upregulated in V1B2+ DCs in both young and old mice, while genes in Q3 are upregulated in V1B2- DCs, also in young and old mice. Quadrants are defined by average log2FC threshold of 0. Avg log2FC=>0.25 and adj P value=<0.05. **h,** Selected pathways from KEGG and GO Biological Processes databases significantly enriched in V1B2- and V1B2+ DCs from young and old mice. Statistical significance was assessed using two-sided Wilcoxon rank sum test (Mann–Whitney *U* test) (b-c, e-h). Only pathways with p adj value <0.05 are shown, enrichment is shown as a combined score, as previously described^56^. Effect sizes were calculated using Cohen’s d and interpreted as negligible (d < 0.2), small (0.2 ≤ d < 0.5), medium (0.5 ≤ d < 0.8), or large (d ≥ 0.8). Only comparisons with medium or larger effect sizes (d ≥ 0.5) are highlighted. \**P* < 0.05, \*\**P* < 0.01, \*\*\**P* < 0.001. Young (*n* = 2) and old (*n* = 2) mice were used (a–h). Gene Ontology (GO), Biological Processes (BP), Cellular component (CC), fold change (FC)

Multi-platform molecular pathway analysis of differentially expressed genes between V1B2+ and V1B2- immune populations in old mice, revealed common patterns of enriched pathways related to lipid metabolism, secretory/vesicle-mediated transport or endocytosis (Fig. 5e). This suggests potential metabolic reprogramming in V1B2+ cells. Additionally, we observed a negative regulation of apoptosis for specific immune cell populations (DCs, AMs) (Fig. 5e), together with an increased Fc-receptor-mediated response in IM and upregulation of phagocytosis/endocytosis-related pathways in AMs, which aligns with macrophage tissue-resident functions. However, V1B2+ cells across all immune populations showed reduced activation of immune responses and lower expression of translation-related genes, consistent with our previous observations in csV1B2+ myeloid population (Fig. 4g). This suggests overall shift towards immune exhaustion of V1B2+ myeloid cells in old mice.

### Myeloid V1B2 expression is associated with immunosuppression

SnCs are heterogenous in nature, and their functional behavior can change over the course of aging^36,38^. We therefore asked if the presence of V1B2+ immune cells with reduced activation of immune responses is an age-dependent, potentially contributing to compromised immune function observed in ageing. To address this question, we performed differential gene expression analysis between V1B2+ immune cells isolated from young and old mice. We focused on AMs, IMs, and DCs populations, with enough V1B2+ cells in both age groups to allow for such analysis. Gene Ontology enrichment analysis of genes upregulated in the V1B2+ immune populations from old mice, revealed gene signatures associated with antigen processing and presentation (Fig. 5f). Interestingly, DCs displayed a decrease in gene signatures related to Fc-receptor mediated signaling, decreased activation of B cell and chemokine signaling and NK cell mediated cytotoxicity, suggesting suppression of adaptive and innate immune responses (Fig. 5f). Similarly, aged V1B2+ IMs, indicated a marked increase in the expression of genes related to regulatory T cell and myeloid differentiation (Fig. 5f). These findings suggest that aging immune cells with csV1B2 reflect a common age-associated shift towards an immunosuppressive phenotype, which could contribute to accumulation of senescent and damaged cells in the aged tissues. However, V1B2+ AMs showed upregulation of genes related to phagocytosis and secretory/vesicle-mediated transport, suggesting that the cell surface expression of V1B2 is associated with the activation of homeostatic functions of alveolar macrophages during ageing (Fig. 5f).

To understand if transcriptional changes of V1B2+ are conserved in ageing at the immune population level, we compared fold change in gene expression between V1B2+ and V1B2- DCs in young and old mice (Fig. 5g). A total of 92 genes were consistently upregulated in V1B2+ DCs in both age groups (Q1), while 84 genes were consistently downregulated (Q3). It indicates that DC induce a robust and age-independent transcriptional signature associated with the membranal presence of V1B2 pump (Fig. 5g). Genes upregulated in V1B2+ dendritic cells (Q1) were primarily enriched in pathways related to monocytes differentiation and negative regulation of apoptosis (Fig. 5h). In contrast, genes upregulated in V1B2- cells (Q3) were associated with activation of immune response. Altogether these results suggest that V1B2+ myeloid immune cell types exhibit an immunosuppressive phenotype that is present irrespective of age, but which is further pronounced in aged tissues.

## Discussion

In this study, we identified V1B2, a subunit of the V-ATPase proton pump, as a cell surface protein enriched on senescent cells. Notably, our findings reveal a novel external orientation for V1B2 anchorage, in contrast to previously documented instances where it is oriented toward the cytosol. Interestingly, only a subset of SnCs exhibits surface expression of V1B2, suggesting its contribution to the heterogeneity of the senescent cell population. Our results suggest that csV1B2 is associated with regulatory mechanism of lysosomal activity and intracellular pH, potentially linking its surface expression to functional diversity among SnCs. In line with this, V1B2+ SnCs show a transcriptional signature associated with DNA damage, repair and resistance to apoptosis. Consistent with this, V1B2+ SnCs are significantly resistant to ABT737-induced senolysis. Overall, our findings suggest that the cell surface expression of V1B2 in SnCs may be part of a selective DNA damage response mechanism that promotes resistance to apoptosis and contributes to their persistence in ageing tissues.

SnCs are heterogeneous population in vivo, with phenotypical variability dependent on the cell type, the microenvironmental niche and the induction mode. This heterogeneity presents a challenge in identifying and targeting SnCs for research and therapeutic purposes. Here, we identified a cell surface expression of V1B2 enriched in cellular senescence that offers a novel targetable marker for the isolation and elimination of apoptosis resistant subset of senescent cells. V1B2 is a subunit of V-ATPase, activity of which is partially conferred through changes in its density on the plasma membrane, which is controlled through reversible exocytosis and endocytosis of v-ATPase containing vesicles^39^. Indeed, V1B2+ cells exhibit significant changes in pathways related to mTOR1 and its counterpart AMPK as well as intracellular trafficking. Moreover, mTORC1 activation suppresses autophagy and promotes the survival of senescent cells. This aligns with our findings of enriched pathways related to autophagy and resistance to apoptosis. It is tempting to speculate that V1B2 membrane expression might promote senescent cell persistent accumulation in ageing and thus might be senescence biomarker correlated with age-related diseases.

Of note, V1B2 can be also expressed extracellularly on non-senescent cells. Thus, while targeting V1B2 could be a promising strategy to eliminate V1B2-positive senescent cells, they could also deplete V1B2-positive non-senescent cells. Therefore, targeting V1B2 in combination with other markers and pathways of SnCs may offer therapeutic opportunities to treat senescence-mediated age-associated diseases.

Senescence is cellular response to stress cues, such as DNA damage, which limits tissue damage in vivo. Thus, SnCs need to be retained in the system to fulfil their physiological role. Interestingly, our results indicate that csV1B2 expression in myeloid cells is associated with DNA damage checkpoints and resistance to apoptosis regardless of age. Thus, we hypotheses that membrane translocation of V1B2 might be the result of the response to DNA damage, which initiates repair mechanisms and anti-apoptosis signaling. Altogether, these results suggest that gene expression program of V1B2+ cells might be associated with specific phenotypic fitness traits such as DNA repair mechanisms in response to DNA damage, regardless of age or microenvironment these cells reside in.

Immunosurveillance of SnCs regulates the dynamics of senescent cell turnover in vivo^28^. Our results suggest that csV1B2 expression on myeloid cells is negatively correlated with activation of immune responses in vivo, as exemplified by enriched pathways related to activation and immune cell trafficking in V1B2- DCs compared to V1B2+ counterparts. Furthermore, expression of PD-L1 or HLA-E on V1B2+ cells suggests that these cells have immunosuppressive phenotype correlated with p16-dependent regulation of PD-L1 stability in SnCs^14^. Consistent with the idea that only a subpopulation of SnCs accumulates with age, V1B2+ SnCs increase in aged tissues and may contribute to immune evasion and prolonged persistence. These properties of V1B2+ cells might also contribute to the role of SnCs in tissue repair and regeneration. Thus, cell surface localization of V1B2 on SnCs may reflect a stress adaptation feature that impairs immune clearance and promotes persistent senescence within tissues.

In conclusion, we have identified that subset of SnCs upregulate V1B2, a subunit of ATPase proton pump, on their cell surface. This localization of V1B2 on SnCs is linked to intrinsic properties of senescent cells in vivo. Our data suggest that SnCs re-localize intracellular V1B2 to the plasma membrane as a response to DNA damage and repair mechanisms, as well as an anti-apoptotic signaling. Immunosuppressive nature of csV1B2+ cells suggests that, in addition to intrinsic resistance to apoptosis, these cells may employ immune evasion strategies to facilitate their accumulation in tissues with age.

Persistence of SnCs in tissues exacerbate age-related symptoms and shortens healthspan and lifespan. Understanding functional heterogeneity of SnCs, marked here by csV1B2 and associated with their turnover in the tissues, might advance the development of more focused, and thus potent, senolytic therapies.

## Materials and Methods

### Cell culture

Human lung fibroblasts IMR-90 (ATCC CCL-186) were cultured in DMEM+10% FBS+1% PenStrept with routine media changes biweekly. Senescence was induced with a 48hr treatment of 100uM Etoposide (Sigma E1383, dissolved in DMSO). Quiescence was induced culturing fully confluent IMR-90 for an additional 12 days with routine media changes. Passaging and harvesting for single cell suspension were performed using Trypsin.

### Mice

C57BL/6 12 months old (“young”) and 22-24 months old (“old”) female mice were purchased from Harlan Laboratories and housed in The Weizmann Institute of Science in accordance with national animal care guidelines. All animal experiments were approved by the institute’s Animal Care and Use Committee.

### Bleomycin intratracheal injections

Mice were anesthetized using a mix of 0.2mg Xylazine (Sedaxylan) and 1mg Ketamin (Ketavet) per 1gr mouse weight. The throat was cut open and 2uL Bleomycin (Sigma B2434) 0.75U/ml per 1gr mouse weight was administered into the trachea with a sharp syringe. The cut was closed with 1-2 clips. 2 weeks post treatment lungs were harvested.

### Lung single cell isolation

To get lung single-cell suspension, mice were euthanized by intraperitoneal injection of ketamin 10 mg/ml (Ketavet Veterinary, 100mg/ml, Zoetis) and xylazine 2 mg/ml (Sedaxylan 20mg/ml, Eurovet) and then transcardially perfused with 10ml cold PBS via the right ventricle. Lung tissues were dissected from mice, cut into small fragments on ice and suspended in 1.5 ml of DMEM/F12 (Invitrogen, 11330-032) containing elastase (3 U ml^−1^, Worthington, LS002279), collagenase type IV (1 mg ml^−1^, Thermo Scientific, 17104019) and DNase I (0.5 mg ml^−1^, Roche, 10104159001) and incubated at 37 °C for 20 min with frequent agitation. After dissociation procedure, cells were washed with an equal volume of DMEM/F12 supplemented with 10% fetal bovine serum and 1% penicillin– streptomycin (Thermo Scientific), filtered through a 100-μm cell strainer and centrifuged at 380*g* for 5 min at 4 °C. Pelleted cells were resuspended in red blood cell ACK lysis buffer (Gibco, A1049201), incubated for 2 min at 25 °C, centrifuged at 380*g* for 5 min at 4 °C and then resuspended in ice-cold fluorescence-activated cell sorting (FACS) buffer (PBS supplemented with 2 mM EDTA, pH 8 and 0.5% bovine serum albumin).

### Flow cytometry

Lung single-cell suspensions were performed by initial staining with live/dead fixable blue commercial kit (L23105, ThermoFisher) according to the manufacturer’s instructions, followed by washing and Fc blocking with CD32 (156604, BioLegend) for 10 minutes. Extra cellular staining was done in BV buffer (00-4409-42, ThermoFisher) with in house anti-V1B2 and secondary antibody (115-605-003, Jackson), CD45 (749889, BD), CD45 (103173, BioLegend), CD31 (102427, BioLegend), EpCam (740281, BD), Ly6c (NB100-65413AF532, Novus), Ly6G (741813, BD), CD11b (101263, BioLegend), CD11c (130-102-797, Miltenyl), CD64 (139311, BioLegend), MHC2 (107645, BioLegend). Fixation and permeabilization was done with a commercial kit (00-5523-00, ThermoFisher) according to the manufacturer’s instructions. Fixed and permeabilized cells were blocked with donkey serum for 10 minutes followed by intracellular staining with p16 (54210, Abcam). All centrifugation steps of live cells were done at 300xg and centrifugation steps of fixed cells were done at 600xg for 5 minutes. Antibodies incubation was performed at 4C for 30 minutes followed by 3 washes with cold PBS.

For analysis of live IMR-90 cells, the cells were stained with DAPI (62248, ThermoFisher) V1B2 (AB73404, Abcam) and secondary antibody (111-605-003, Jackson). Cells were analyzed with CytoFLEX flow cytometer.

### Imaging Flow Cytometry

Cell were imaged by Imaging flow cytometer (ImageStreamX Mark II, AMNIS corp.–part of Cytek, CA, USA). The cells were imaged using a 40X lens (NA=0.75). Lasers intensities were set to maximum power that did not cause pixel saturation. The lasers used were 405nm, 488nm, 561nm, 642nm and for side scatter collection 785nm (2mW). The channels acquired were Brightfield (Ch01, Ch09), V1B2 antibody (Ch02 or Ch11), side scatter (Ch06).

IMR-90 cells were stained with zombie NIR (77184, BioLegend) for live/dead discrimination, andti-hV1B2 and secondary antibody (109-545-044, Jackson) followed by fixation and permeabilization as described above and intracellular staining with V1B2 (AB73404, Abcam) or cathepsinB (EPR4323, Abcam) and secondary antibody (111-605- 003, Jackson). For the lung tissue experiment the additional channels used were Ch04 (CD11c), Ch07 (SiglecF), Ch10 (CD45), and Ch11 (p16).

Data was analyzed using IDEAS 6.3 (Amnis Corp.). Cells were gated either for DNA+ or negative for the live/dead stain, vs. brightfield area. Focused cells were further selected by the Gradient RMS and Contrast features (measuring changes in pixel intensities within the image). Cropped cells were eliminated by plotting the Centroid X (The distance from the left side of the screen) vs. Area of the brightfield image. For figure 1d, cells were further gated for more circular cells by using the Circularity feature (The degree of the mask’s deviation from a circle), vs. aspect ratio of the bright-field image. To further filter out debris and damaged cells, we gated on the bright field vs. nuclear area. V1B2 staining was quantified by using the intensity and max pixel values of the V1B2 channels.

Membrane staining was calculated by using the combined mask of the bright field (M01) excluding the 80% eroded mask of the image (M01 And Not AdaptiveErode (Object(M01, BF, Tight), BF, 80).

The bGal gating was done according to the bright-field mean pixel values, as described in Biran et al^40,41^.

### pH measurements

BCECF, AM (ThermoFisher) 20uM for IMR-90 (analyzed in CytoFlex) and 0.2uM (analyzed in Aurora conventional mode) was added to the antibody staining step in a flow cytometry protocol. A calibration curve was created as per product guidelines using pH buffers (ThermoFisher) 10μM Nigrecin and 10μM Valinomycin.

### ABT-737

For mice IP injections ABT-737 was dissolved in a solution of 3.25% Dextrose, 30% propylene glycol, 5% Tween 80 and administered in two consecutive daily injections of 25mg/kg. 48hr post end of treatment mice were sacrificed. For IMR-90 tissue culture treatment media was supplemented with 10uM ABT-737 dissolved in DMSO and harvested the following day for analysis.

### X-gal staining

Harvested cells were fixed with 4% PFA for 5 min while vortexing; then washed twice for 5 min in 1mM MgCl2 in PBS pH 5.5 for mouse pH 6 for human and incubated for 8hr at 37^0^C with x-gal staining solution (1mg/ml x-gal (Roche), 5uM K_3_Fe(CN)_6_, 5uM K_3_Fe(CN)_6_, 1mM MgCl_2_ in PBS). Cells were washed x2 with PBS and then fixed and permeabilized with a commercial kit by ebiosciences 30 min at 4C in the dark. Staining procedure followed as specified in flow cytometry section.

### Bulk RNAseq and analysis

Sequencing libraries were prepared using MARS-seq protocol^42^. Single-end reads were sequenced on 1 lane of an Illumina NextSeq. The output was ∼11 million reads per sample.

Bulk MARS-Seq was analyzed using the UTAP pipeline^43^. Poly-A/T stretches, and Illumina adapters were trimmed from the reads using cutadapt^44^; resulting reads shorter than 25bp were discarded. Remaining reads were mapped onto 3’ UTR regions (1,000 bases upstream of the 3’ end and 100 bases downstream) of the H. sapiens, hg38 genome according to RefSeq annotations, using STAR^45^ v2.4.2a with EndToEnd option and outFilterMismatchNoverLmax was set to 0.05, twopassMode Basic and alignSoftClipAtReferenceEnds set to No. Deduplication was carried out by flagging all reads that were mapped to the same gene and had the same UMI. Counts for each gene were quantified using htseq-count^46^ in union mode, using the gtf above and corrected for UMI saturation. Differentially expressed genes were identified using DESeq2^47^ with the betaPrior set to True, cooksCutoff and independentFiltering parameters set to False using batch correction with the “sva” package. Raw P values were adjusted for multiple testing using the procedure of Benjamini and Hochberg. Significant differentially expressed genes were determined by values of padj <= 0.05, log2FoldChange| >= 1 and baseMean >= 5. Pipeline was run using snakemake^48^. Pathway analysis and enrichment was run using GeneAnalytics tool^49^.

### FACS sorting for single cell RNA sequencing

Lung single-cell suspension was stained with anti-mouse CD16/32 (1:200, eBioscience, 14-0161-82) to block Fc receptors, followed by staining with CD45-BV605 (103-139, Biolegend) and APT6V1B2 (ab73404). To exclude live/dead and endothelial cells, cells were stained with Sytox Blue (Invitrogen, 34857) and CD31-Pacific Blue (Biolegend, 102421) respectively.

To allow sample multiplexing for single-cell transcriptomics, we used Cellular Indexing of Transcriptomes and Epitopes by Sequencing (CITE-Seq). CITE-Seq enables combined single-cell RNA sequencing, cell surface marker detection using antibody-derived tags^50^, and labeling of individual samples using antibody “hashtags”^51^. For CITE-seq, cells were stained with Biolegend TotalSeq™ B oligo-conjugated antibodies for 30 minutes at 4°C, washed thoroughly, and resuspended in PBS + 0.04% BSA. Samples were labelled with *hashtag* antibodies as follows: Young 1 (Epcam *118247; CD45, 155843),* Young 2 (*CD45, 155845), Old 1 (*Epcam, *118247; CD45, 155847), Old 2 (CD45, 155849)*.

Before sorting, all samples were filtered through a 70-μm cell strainer. Dead cells, endothelial cells and doublets were excluded by gating, after which CD45+ cells were gated based on BV605 fluorescence, followed by gating for the ATP6V1B2 staining. 10,000 - 20,000 CD45+ immune cells from young and old mice were sorted based on their V1B2 membrane expression into a pre-cooled 1,5 mL eppendorf tube with 0.04% BSA in PBS using a BD FACS Aria Fusion flow cytometer (BD Biosciences).

### single cell RNA-sequencing and analysis

#### Library preparation and sequencing

Single cell RNA-sequencing (scRNA-seq) libraries were prepared using the chromium single cell RNA-seq platform (10x genomics) according to manufacturer’s instructions. 30,000 single-cell suspension was loaded onto Next GEM Chip G targeting 15,000 cells and then ran on a Chromium Controller instrument to generate GEM emulsion. Single-cell 3’ RNA-seq libraries and cell surface protein libraries were generated according to the manufacturer’s protocol (Chromium Single Cell 3’ Reagent Kits User Guide (v4 Chemistry Dual Index). Final libraries were quantified using NEBNext Library Quant Kit for Illumina (NEB) and high sensitivity D5000/D1000 TapeStation (Agilent). Libraries were pooled according to their molarity, aiming for ∼25,000 reads per cell for gene expression libraries and ∼5,000 reads per cell for cell surface protein libraries. Pooled libraries were sequenced on a NovaSeq X sequencer using and 100 cycles reagent kit (Illumina) with 100 cycles with a depth of 1.5 billion reads per sample.

#### Pre-processing and quality control

Data was pre-processed with the 10x Genomics CellRanger software (v. 8.0.1) and further annotated on the mm10 reference set (refdata-gex-mm10-2020-A provided by 10x Genomics). Demultiplexing with hashtag oligos (HTOs) was performed as previosuly described^51^. Cells labeled with two hashtag oligos originating from different samples were removed. Quality control, clustering and downstream analysis was performed using R (v. 4.3.1) (R Core Team (2020) using Seurat package (v. 5.0.0). Briefly, doublets and multiplets were eliminated based on gene counts > 4445 and UMI counts > 17489. Putative dead Cells and empty droplets were discarded based on mitochondrial gene percent > 4.2, gene count < 445 and UMI count < 696. This resulted in a dataset of 53,432 single cells. Young and old mouse samples amounted to 32174 and 21258, respectively.

#### Clustering

Normalization and scaling were carried out separately on each sample using the *SCTransform* function in Seurat. The top 3,000 highly variable genes were identified using *SelectIntegrationFeatures* and subsequently merged into one object using the Seurat *merge* function. Cells were clustered using the *FindNeighbors and FindClusters* functions with 17 principal components and a resolution parameter of 1.2. Cells were visualized in two-dimensional space using the Seurat *RunUMAP* function which implements Uniform Manifold Approximation and Projection (UMAP) for dimension reduction. Cell cluster identities were determined based on the expression of canonical marker genes using the *FindAllMarkers* or *FindMarkers* functions with default parameters.

#### K-means clustering

To identify subgroups of genes with similar patterns of regulation across conditions, k-means clustering analysis was performed using the *kmeans* function on the average expression of all genes that were differentially expressed in at least one comparison between conditions. A heatmap was constructed from data generated using k=4 and the R package ggplot.

#### Differential gene expression analysis

Differential gene expression analysis was performed using the Seurat package and Wilcoxon Rank Sum test between experimental conditions. Genes were determined to be differentially expressed based on a threshold of log2 fold change >= 0.25 and FDR (Benjamini-Hochberg) -adjusted p value <= 0.05 and, visualized as a volcano plot (Enhanced Volcano R package)^52^. For heatmap visualization of differentially expressed genes, data that was scaled (z-score) using the Seurat *SCTransform* function was plotted using the Seurat *DoHeatmap* function^53,54^).

Furthermore, average gene expression per cluster was calculated using the Seurat AverageExpression function on log-normalized scaled data. Senescence and apoptosis resistance scores were calculated based on normalized expression of gene sets associated with cellular senescence ^35^ and apoptosis resistance (Bcl2, Bcl2a1b, Bcl2a1d, Rb1, Birc6, Xiap), respectively. Function AddModuleScore was used in order to create a single feature score for each list.

GO and pathway enrichment analysis was performed in k-means clusters (all genes), general myeloid population and identified immune cell populations (up to 100 top differentially expressed genes) with the GO Biological Process 2023, GO Cellular Component 2023, GO Molecular Function 2023, KEGG 2019 Mouse, WikiPathways 2024 Mouse, Elsevier Pathway Collection databases using the Enrichr R package^55^ The significance of the tests was assessed using a combined score, described as *c* = log(*p*) × *z*, in which *c* is the combined score, *p* is the Fisher’s exact test *P* value and *z* is the *z*- score for deviation from expected rank^56^.

## Acknowledgements

We thank Y. Berkovich for help with animal procedures, V.K. was supported by grants from the European Research Council (856487), from the Israel Science Foundation (1626/20), DFG - CRC 1506 “Aging at Interfaces”, Weizmann - Nella and Leon Benoziyo Center for Neurological Diseases, Weizmann - Sagol Centers for Research on the Aging Brain and for Longevity Research, Weizmann SABRA - Yeda-Sela - WRC Program, the Estate of Emile Mimran, and The the Maurice and Vivienne Wohl Biology Endowment, Shimon and Golde Picker Award, the Estate of Gerald Alexander and Moross Integrated Cancer Center V.K. is the Director of EKARD Institute for Cancer Diagnosis Research.

## Authors contribution

Conceptualization of the study was carried out by N.F., J.M., and V.K. The methodology and investigation were planned and executed by N.F., J.M and Y.O. Ideation was contributed by N.F., J.M., E. K., Z.P., A.M., T.S and G.S. Visualization was performed by N.F., J.M., A.M., and B.D. Supervision was undertaken by V.K. and U.A. Writing of the original draft was performed by N.F. and J.M. Review and editing was carried out by N.F., J.M. and V.K. Funding acquisition and project supervision was performed by V.K.

## Competing interests

N.F., Y.O. and V.K. are co-inventors on patent applications related to the topic of this study. The other authors declare no competing interests.

## Data availability

All NGS sequencing data in this manuscript are available at NCBI GEO under the accession numbers GSE390197 (Bulk RNA-seq data of V1B2+ and V1B2- cells in damage-induced senescence) and GSE390170 (single-cell RNA-seq data of V1B2+ and V1B2- lung immune cells).

---

**Extended Data Figure 4.**
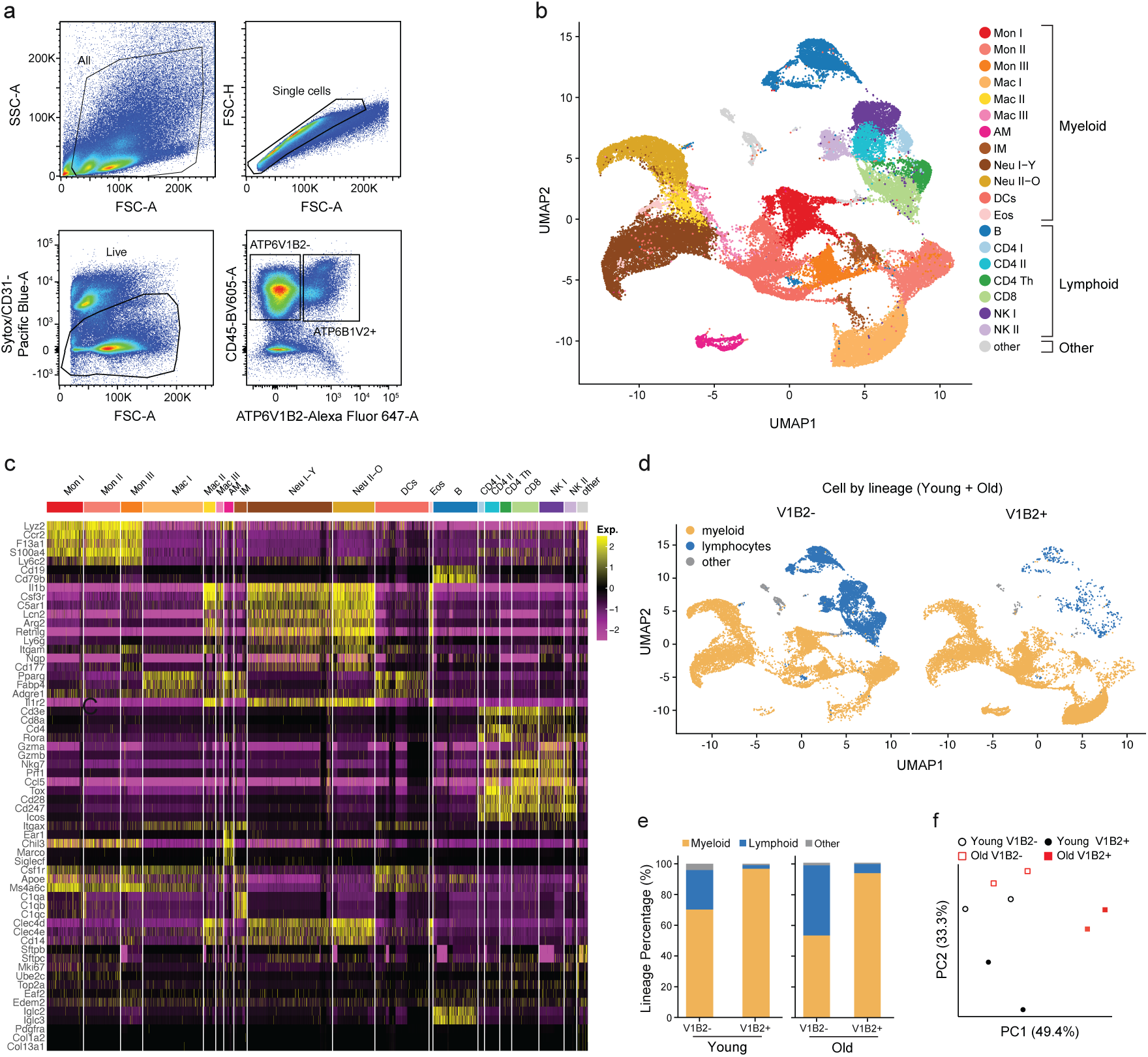
Myeloid-biased V1B2 expression is linked to an age-independent response to DNA damage. **a,** Gating strategy for sorting V1B2- and V1B2+ CD45⁺ cells by FACS. **b,** Uniform manifold approximation and projection (UMAP) visualization of 50 000 single cells from lung samples of young and old mice. Color represents different immune cell-types obtained by clustering and annotated by conventional markers. **c,** Heatmap showing expression of canonical marker genes used to define immune cell populations. **d,** UMAP visualization of immune cell lineages within V1B2- and V1B2+ cells. Cells were down sampled to 16518 cells. **e,** Bar plots showing the distribution of immune lineages in V1B2- and V1B2+ cells in young (left) and old (right) mice. **f,** Principal components (PC) analysis based on transcriptomic profiles of V1B2-negative and V1B2-positive myeloid cells from young and old mice, with two biological replicates per condition.

**Extended Data Figure 5.**
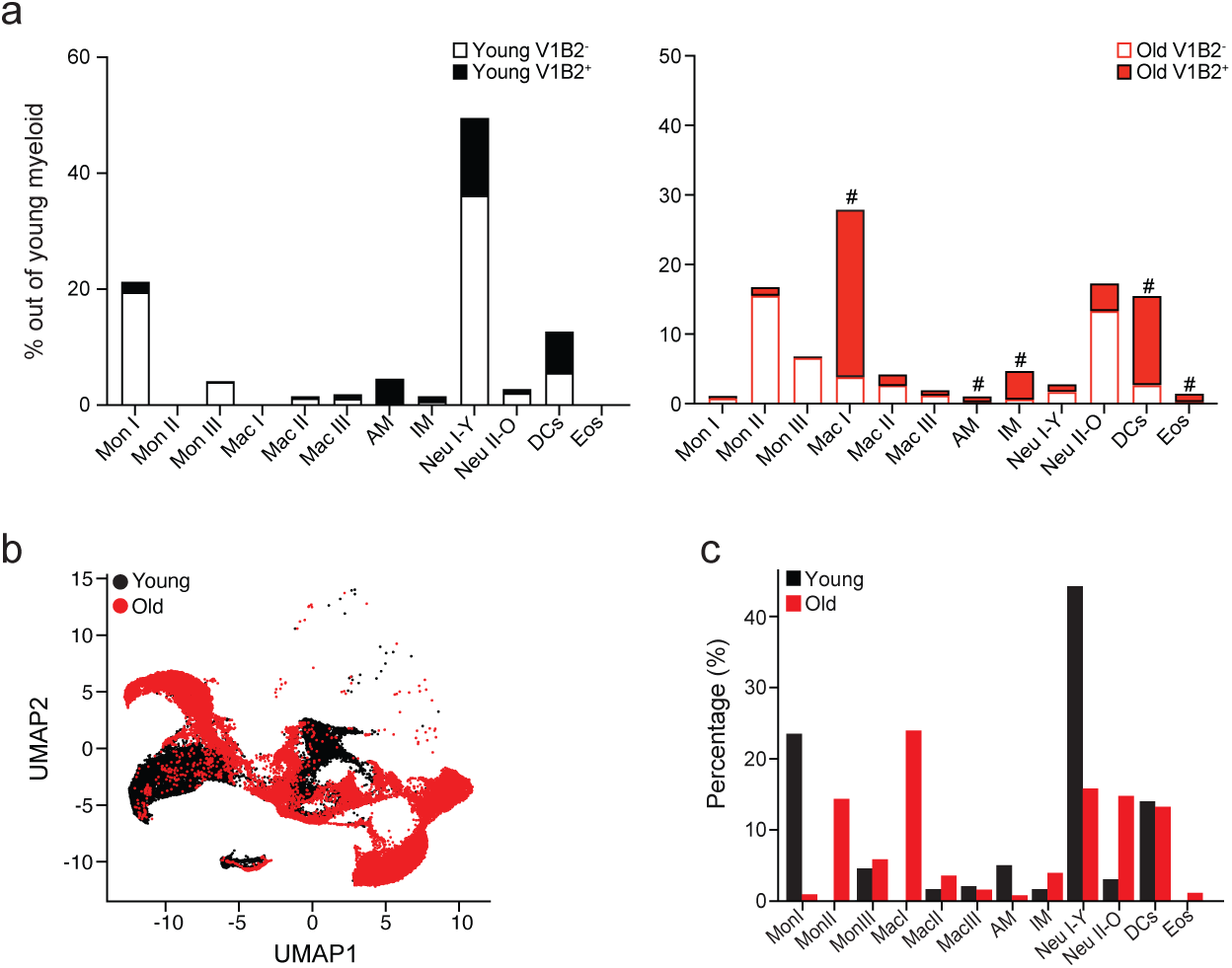
V1B2 expression is associated with senescence and resistance to apoptosis *in vivo*. **a,** Distribution of V1B2+ and V1B2- immune cell types in young (left) and old (right) mice. Immune cell types with enrichment of V1B2+ cells are highlighted. **b,** UMAP of myeloid lineage from young and old mice. **c,** Bar plot showing percentage of identified myeloid immune cell types in young and old mice.

